# Selection favors adaptive plasticity in a long-term reciprocal transplant experiment

**DOI:** 10.1101/2020.08.31.275073

**Authors:** Jill Anderson, M. Inam Jameel, Monica A. Geber

## Abstract

Spatial and temporal environmental variation can favor the evolution of adaptive phenotypic plasticity, such that genotypes alter their phenotypes in response to local conditions to maintain fitness across heterogeneous landscapes. When individuals show greater fitness in one habitat than another, asymmetric migration can restrict adaptive responses to selection in the lower quality environment. In these cases, selection is predicted to favor traits that enhance fitness in the higher-quality source habitat at the expense of fitness in the marginal habitat, resulting in specialization to the high-quality environment. Here, we test whether plasticity is adaptive in a system regulated by demographic source-sink dynamics. *Vaccinium elliottii* (Ericaceae) occurs in dry upland and flood-prone bottomland forests throughout the southeastern United States, and shows patterns consistent with source-sink dynamics. We conducted a multi-year field experiment to evaluate whether plasticity in foliar morphology is advantageous. Both across habitats and within the high-quality upland environment, selection favored plasticity in specific leaf area and stomatal density. Stabilizing selection acted on plasticity in these traits, suggesting that extreme levels of plasticity are disadvantageous. We conclude that even in systems driven by source-sink dynamics, temporal and spatial variation in conditions can favor the evolution of plasticity.

## Introduction

Species that inhabit spatially or temporally heterogeneous landscapes often exhibit phenotypic plasticity, such that individuals shift their phenotype in response to environmental stimuli (e.g., Dudley and Schmitt 1996; Boersma et al. 1998; Galloway and Etterson 2007; Lind and Johansson 2007; Forsman 2015; Hendry 2015). If individuals can sense and react to reliable cues, selection can favor plasticity under temporal variation, when individuals experience multiple conditions across their lifespan (Moran 1992; Stratton and Bennington 1998), and under spatial variation when progeny establish in non-parental habitats (Alpert and Simms 2002). For example, populations of the annual plant *Erodium cicutarium* maintain higher plasticity in spatially heterogeneous serpentine soil patches than in the more homogeneous non-serpentine areas (Baythavong 2011). Additionally, adaptive plasticity can enable population persistence during environmental change (Charmantier et al. 2008; Nicotra et al. 2010). However, phenotypic plasticity could also be neutral or represent a maladaptive or passive response to stress (Hendry 2015). Evaluating the fitness consequences of plasticity is crucial for predicting evolutionary responses to environmental heterogeneity (Nicotra et al. 2010). Nevertheless, testing whether plasticity confers a fitness advantage remains challenging because analyses require fitness and trait data from replicated accessions transplanted into at least two environments, ideally in natural habitats in the field.

Plasticity could be especially advantageous for species with spatially-extensive gene flow because offspring can disperse broadly into different environments (Alpert and Simms 2002; Hendry 2015). To that point, the amount of plasticity in island populations of the frog, *Rana temporaria*, increased as a function of the amount of gene flow from populations in disparate habitats, along with the degree of local environmental variation (Lind et al. 2011). In addition, plasticity could enhance fitness for long-lived species, which experience multiple years of fluctuating conditions before reaching reproductive maturity (Bradshaw 1965). For example, directional selection favored morphological plasticity in response to flooding and competition in a clonal perennial buttercup (*Ranunculus reptans*) (Van Kleunen et al. 2007). We hypothesize that stabilizing selection could also operate on trait plasticity. Stabilizing selection often favors intermediate phenotypes (e.g., Dudley 1996; Brooks et al. 2005; Wadgymar et al. 2017; Taylor et al. 2018), but few studies have evaluated nonlinear selection on trait plasticity. Canalized genotypes with limited plasticity could be at a fitness disadvantage under spatial or temporal variation because they cannot shift their phenotypes. Similarly, highly plastic lines could also experience reduced fitness if they are too phenotypically labile, either altering phenotypes too readily in response to environmental variation or expressing exaggerated trait values. Thus, we might expect fitness to be maximized at an intermediate trait plasticity.

Many species inhabit landscapes in which habitat patches vary in quality or some habitat types occur more frequently (Kawecki 2008). The evolution of adaptive plasticity could be constrained if habitat quality differs, such that individuals have higher fitness in some habitat than others, or if habitat types vary in abundance. In the case of demographic source-sink dynamics, migration from source populations sustains sink populations; this asymmetric migration could potentially counteract selection within sink populations, leading to local maladaptation there (Pulliam 1988; Sultan and Spencer 2002; Kawecki 2008). In these systems, traits favorable in the source environment are expected to evolve at the cost of adaptations to the marginal habitat (Kawecki 2008). Here, we extend this logic to the evolution of plasticity.

Adaptive phenotypic plasticity is a strategy that maximizes fitness across habitat types (Baythavong and Stanton 2010; Baythavong 2011). In systems regulated by source-sink dynamics, the evolutionary response to selection is biased toward traits that are adaptive in the more frequent or higher quality source environment (Holt and Gaines 1992; Stanton and Thiede 2005; Kawecki 2008). Given the potential costs and limitation of plasticity (DeWitt et al. 1998), we would not expect adaptive plasticity to evolve in response to conditions in the sink environment under source-sink population dynamics unless selection within the source habitat favors plasticity. Instead asymmetrical gene flow in a source-sink system could result in the evolution of specialization to the source environment (Holt and Gaines 1992; Sultan and Spencer 2002).

The high bush blueberry, *Vaccinium elliottii* (Ericaceae), is a perennial woody shrub endemic to the southeastern United States, where it grows across a gradient of water stress from seasonally flooded bottomland hardwood forests with dense canopies to more arid upland forests with high light levels in the understory (Radford et al. 1968; Godfrey and Wooten 1981; Anderson et al. 2010). These contrasting conditions could impose divergent natural selection, favoring alternate phenotypic optima in each habitat. This species demonstrates demographic source-sink dynamics (Pulliam 1988), as reciprocal transplant experiments and genotyping via microsatellite markers suggest that asymmetric gene flow from abundant upland populations into sparse bottomland populations could constrain adaptation to bottomland forests (Anderson and Geber 2010). Nevertheless, *V*. *elliottii* expresses extensive plasticity in morphology (specific leaf area, foliar nitrogen content, root:shoot ratio, allocation to shallow roots) and physiology (photosynthesis, stomatal conductance and water use efficiency) to flood vs. drought treatments in the greenhouse, and to bottomland vs. upland forests in the field (Anderson et al. 2010). Thus, this system presents a disconnect between the expectation that selection should favor adaptations to the source environment (upland habitats) and the observation of extensive plasticity across habitat types.

Here, we examine selection on plasticity in three foliar traits (Table 1), which are linked to physiological function and subject to divergent selection across flooding/aridity gradients in other systems: specific leaf area, leaf lamina area (hereafter: leaf area), and stomatal density (Steinger et al. 2003; Wright et al. 2004; Carlson et al. 2015; Maire et al. 2015; Ramírez-Valiente et al. 2018). Stomatal anatomy influences the rate of stomatal conductance (Lawson et al. 1998; Franks and Beerling 2009). A recent meta-analysis revealed that stomatal density increases with light intensity across species (Poorter et al. 2019), which leads to the hypothesis that selection would favor increased stomatal densities in the high-light upland environment. Alternatively, selection could favor lower stomatal density in arid upland environments to prevent water loss from transpiration (Woodward et al. 2002; Carlson et al. 2015). Specific leaf area is often correlated with photosynthetic rate and typically decreases in high-light and arid environments (Steinger et al. 2003; Wright et al. 2004; Terashima et al. 2011; Maire et al. 2015), leading to our prediction that selection favors reduced specific leaf area in upland habitats. Finally, arid, high light environments induce small leaves in other systems (Valladares et al. 2000; Carlson et al. 2015; Ramírez-Valiente et al. 2018), and we predict that selection in upland environments will favor reduced leaf area.

**Table 1:** Predictions of trait variation across habitat types, divergent selection, and selection on plasticity. For each trait, we indicate whether data from this study support the predictions and reference the corresponding figure. *Vaccinium ellioittii* achieves greatest fitness in upland environments.

|  |  | **Specific Leaf Area (SLA)** | **Stomatal density** | **Leaf area** |
| --- | --- | --- | --- | --- |
| **Trait plasticity** | **Predictions** | Higher in bottomlands than uplands | Higher in uplands than bottomlands | Larger in bottomlands than uplands |
|  | **Results** | Supported for all years (Fig. 1a, 1d) | Supported for some years (Fig. 1c, 1f) | Not supported (Fig. 1b, 1e) |
| **Divergent selection** | **Predictions** | Selection for higher SLA in bottomlands than in uplands | Selection for reduced stomatal density in bottomland than in uplands | Selection for larger leaves in bottomlands than in uplands |
|  | **Results** | Supported (Fig. 2c) | Not supported (Fig. 2a) | Not supported; instead stabilizing selection favored intermediate leaf area (Fig. 2b) |
| **Selection on plasticity** | **Predictions** | Selection favors plasticity | Selection favors plasticity | Selection favors plasticity |
|  | **Global analysis** | Supported: Stabilizing selection for intermediate plasticity (Fig. 3a) | Supported: Selection for increased plasticity in specific leaf area (Figs. 3b and c). | Not supported: No pattern |
|  | **Within uplands only** | Supported for both cohorts (Fig. 4c, d) | Supported: Directional selection for increased plasticity for 2005 cohort (Fig. 4a) and stabilizing selection for 2006 cohort (Fig. 4b) | Not supported: No pattern |

We test the hypothesis that phenotypic plasticity is adaptive by (1) examining how trait values vary with transplant habitat and growing season to quantify spatial and temporal plasticity; (2) investigating whether divergent selection across habitat types accords with the direction of plasticity and (3) determining whether plasticity confers a fitness advantage across the landscape. For example, if the bottomland environment induces higher trait values than upland forests (as is the case for specific leaf area), we predict that selection should favor a larger trait optimum in the bottomlands and a smaller trait optimum in the uplands, and that plasticity in this trait should be associated with greater fitness averaged across habitat types. If temporal or spatial variation within the source environment favors plasticity within that habitat type, then adaptive plasticity could evolve across the landscape despite source-sink population dynamics.

For this reason, we also hypothesize that plasticity is beneficial within the source (upland) habitat. Finally, we assess nonlinear selection to test whether stabilizing select favors intermediate levels of plasticity. To evaluate our hypotheses, we leverage data from a multi-year field experiment exposing individuals of a woody perennial plant to the suite of environmental factors that differ between discrete habitat types.

## Methods

### Focal system

*Vaccinium elliottii* (Ericaceae, Elliott’s blueberry) is an outcrossing highbush blueberry, which produces insect-pollinated flowers in March-April and sets animal-dispersed seeds in June-July (Martin et al. 1951; Anderson and Geber 2010). This species has low population genetic differentiation (F_ST_= 0.032) and high rates of gene flow between populations within and across habitat types (Anderson and Geber 2010). We conducted fieldwork in the Coastal Plain of South Carolina, where *V*. *elliottii* inhabits xeric upland and flood-prone bottomland forests. We established reciprocal field gardens in two upland and two bottomland forest sites at Francis Beidler Forest, a National Audubon Sanctuary in the diffuse brown-water floodplain of Four Holes Swamp (33° 13’N 80° 20’W) (Anderson et al. 2010). We sampled natural populations throughout the Four Holes Swamp watershed and in the Pee Dee and Santee watersheds of S.C., all of which share similar climates (Anderson et al. 2010). In these systems, bottomland hardwood forests flood 3-139 days/year (average ± SD: 43.6 ± 26.1 days/year), but floodwaters are typically no deeper than several centimeters during a flooding event (Anderson et al. 2010). During the growing season, precipitation ranges from 0-377 mm/month (average ± SD: 125.3 ± 79.8 mm/month), which can induce drought stress in upland forests (Anderson et al. 2010). The clay soils of bottomland forest are nutrient rich relative to sandy upland forest soils, but the dense canopy restricts light to the understory (Anderson et al. 2010).

Historically, upland forests dominated the landscape of the southeastern U.S., and these forests were dissected by river systems associated with large tracts of wetland forests (Hickman 1990; Phillips 1994, and references therein). Across this region, human activities have caused extensive loss of forested habitat (Abernethy and Turner 1987; Hickman 1990; Carter and Biagas 2007; Cubbage et al. 2018). Humans disproportionately converted upland forests to agriculture owing to favorable drainage conditions (Phillips 1994). Cubbage et al. (2018) estimated that bottomland hardwood forests (not permanently flooded swamps) cover ∼9.3 million hectares in 13 states of the southeastern USA (∼11.4% of all timberland in this region), whereas non-wetland upland forests cover ∼47.8 million hectares (∼58% of timberland). Thus, despite large-scale deforestation, upland forests still occur with greater frequency than bottomland hardwood forests in the southeastern U.S.

Previous work with *V*. *elliottii* documented plasticity in morphological and ecophysiological traits, asymmetric gene flow from upland to bottomland, and demographic source-sink dynamics with transplants from both habitat types expressing higher survival and growth in upland than bottomland forests and under drought than flooded conditions (Anderson and Geber 2010; Anderson et al. 2010). Additionally, Anderson and Geber (2010) found that the abundance of adult *V*. *elliottii* individuals was five times greater in upland than bottomland populations, and naturally-occurring upland individuals had >13.5 times greater reproductive success than their bottomland counterparts. Here, we expand upon this earlier examination of patterns of plasticity in specific leaf area by quantifying the extent of spatio-temporal plasticity in three foliar traits (specific leaf area, stomatal density, and leaf area) across a longer timeframe and examining divergent selection on traits as well as selection on plasticity in these traits. Specifically, we conducted genotypic selection analyses, including field fitness from planting (2005-2006) through April 2014 and trait data measured during 2-3 growing seasons to provide the long-term records necessary for evaluating selection in this perennial species. We find relatively low levels of correlations across traits in this experiment (Table S1).

This study focused on vegetative cuttings taken from adult plants in the field that had experienced multiple episodes of selection across their lifespans. To propagate adult tissue, we collected 2-5 cuttings (10cm of new growth) from 20-30 adult plants in 17 upland and 15 bottomland populations throughout South Carolina in the summers of 2004 and 2005 (Anderson et al. 2010). We stored cuttings on ice in the field. In the greenhouse, we applied rooting hormone to the stem (Rhizopon AA #3, 0.8% IBA, Rhizopon bv, Hazerswoude, Holland), and positioned cuttings under an automated misting system for 2-3 months until roots established. We grew rooted cuttings in the greenhouse until May (2005 and 2006) when they were ∼20 cm tall and had woody tissue.

Typically, researchers rear field-collected seeds under greenhouse conditions for a generation to homogenize maternal effects prior to conducting common garden experiments. However, that procedure is not possible when the focal species is a long-lived woody plant that takes many years to reproduce. We minimized variation in maternal effects by growing plants in the same environment under benign greenhouse conditions for 6 months prior to the initiation of the reciprocal transplant experiments. If maternal effects were prominent in our system, we would have expected experimental transplants to show patterns that resemble local adaptation (Galloway and Etterson 2007). Instead, transplants had elevated fitness in upland transplant gardens and depressed fitness in bottomland gardens (Anderson and Geber 2010), suggesting that maternal effects are minimal.

### Field reciprocal transplant

In spring 2005, we transplanted N=1685 cuttings from 412 genotypes and 22 populations into two upland and two bottomland common gardens in the Four Holes Swamp. We expanded this study in spring 2006, when we transplanted N=548 cuttings (106 genotypes from 22 populations) into these same experimental gardens. Some families were represented by only one individual within a transplant habitat, precluding genotypic selection analysis. We restricted the dataset to families for which at least two individuals were planted into each transplant habitat, resulting in a sample size of N=1189 cuttings in the 2005 cohort (mean ± SD: 3.23 ± 1.45 individuals per family per habitat type; 183 genotypes; 17 source populations), and N=466 cuttings in the 2006 cohort (2.94 ± 0.97 individuals per family per habitat type; 79 genotypes; 13 source populations).

To reduce transplant shock, we watered all experimental individuals two times per week for two weeks after planting. Flooding stress differed substantially between the two transplant years, with growing season rainfall exceeding the long-term average by 51mm/month in 2005 (N. Brunswig and M. Dawson, unpub. precipitation records). Additionally, after approximately half of the bottomland transplants were established in 2005, a 45-day long flood occurred in bottomland sites, and the water table remained high even after the floods receded (Anderson and Geber 2010; Anderson et al. 2010). In contrast, monthly precipitation was 16 mm lower than the average growing season value during the 2006 season. By replicating this field experiment, we captured temporal environmental variation in conditions during establishment.

We monitored experimental individuals from 2008 (the last sampling point included in Anderson and Geber 2010; Anderson et al. 2010) until 2014. During the first two years of growth, we visited each individual twice per month to record the time of mortality. Subsequently, we visited each plant in October 2007, March 2008, March 2009, April 2011, October 2012, March 2013 and April 2014. In October of 2006, 2007 and 2012, we collected an average of 5 living sun and 5 living shade leaves per living plant, scanned leaves to extract leaf areas with ImageJ (Schneider et al. 2012), dried leaves at 50°C for 3-4 days, and weighed them on a Mettler AE 200 balance (± 0.0001g) to determine specific leaf area (leaf area per unit biomass, cm^2^/g). On these understory shrubs, we collected both sun and shade leaves to quantify individual-level foliar traits more accurately and precisely. Some leaves had evidence of herbivory. To obtain the leaf area of undamaged leaves (without herbivory), we filled in internal holes in Image J and redrew leaf margins. Owing to low herbivore damage (mean ± SD: 2.1% ± 3.4% leaf area removed by herbivores in N=372 genotypes of the 2005 cohort and 3.1% ± 4.2%, N=298 genotypes in the 2006 cohort), it was straightforward to modify the leaf images to reflect leaf size prior to herbivory.

We quantified stomatal density by making epidermal impressions of the abaxial (lower) leaf surface with clear nail polish, mounting these impressions on microscope slides, and visualizing them under 400 × magnification using a compound microscope. We calculated stomatal density by averaging the number of stomata across four distinct nonoverlapping 0.0352 mm^2^ areas of each impression. Additionally, we made stomatal peels of N=33 samples on the adaxial (upper) surface of the leaf to examine the potential for adaxial stomata (Woodward 1986), which are typically rare in shrubs (Muir 2015). As we were unable to detect any evidence of adaxial stomata in *V*. *elliottii*, we proceeded with quantification of abaxial stomatal density.

## Statistical analyses

For all analyses, we first calculated family-mean trait values for each year of measurement as well as across all years (least square means; hereafter: LSMEANs) and fitness components as a function of transplant habitat by family in models that included block nested within transplant site as a random effect (Proc Mixed, SAS ver. 9.4). We standardized traits to a mean of zero and a standard deviation of one to facilitate comparison of selection on traits measured on different scales. We analyzed the two cohorts separately because of differences in the duration of the monitoring.

### Phenotypic plasticity

We evaluated plasticity across habitat types through a repeated measures multivariate regression with a Kenward-Roger degree of freedom approximation. We analyzed family-level LSMEANs in all three foliar traits jointly as a function of transplant habitat type, year of measurement, source habitat, and all two and three-way interactions, with random effects for family and family by transplant habitat using the Mixed procedure in SAS (ver. 9.4). These multivariate repeated measures models specify the covariance structure of the R matrix using direct (Kronecker) product structures [type=UN@AR(1)] to fit multiple response variables (unstructured covariance matrix, UN) measured on the same plant genotypes across years [autoregressive covariance matrix, AR(1)] (Galecki 1994). A significant main effect of transplant habitat would indicate spatial plasticity, and a main effect of growing season would point to temporal plasticity. Interactions of transplant habitat and season would suggest that the degree of spatial plasticity depended upon the growing season. A main effect of source habitat or interactions with that factor would suggest genetic differentiation in phenotypes between upland and bottomland source populations.

### Genotypic selection analyses

Genotypic selection analyses (Rausher 1992) tested whether: 1) divergent selection favors different phenotypic optima under contrasting environmental conditions, and 2) phenotypic plasticity in foliar traits is adaptive. Many individuals died before foliar traits were measured. These individuals could have died due, at least in part, to limited phenotypic plasticity and trait values that were inappropriate for the transplant environment. Phenotypic traits of dead individuals can be estimated based on trait values of their surviving relatives (Hadfield 2008). Thus, for each family, we calculated genotypic mean fitness based on data from every planted individual, and genotypic mean trait values from individuals that survived until trait measurement.

As very few individuals successfully flowered in the 8-9 years of this field experiment, we focused on survival as a critical component of fitness. For each individual plant, we calculated longevity as the number of elapsed days between planting and mortality. At the final census in April 2014, 479 individuals (40.2%) of the 2005 cohort and 195 individuals (42%) of the 2006 cohort remained alive. In the terminology of survivorship analysis, these individuals would be considered right-censored as they had not yet experienced mortality. However, it is not possible to analyze selection on plasticity within the framework of a survivorship analysis like Cox proportional hazards models because the genotypic selection analyses require family-level data on plasticity whereas survivorship models require individual-level data. Therefore, to include individuals that were alive on the final census in our analyses, we assigned them time of mortality of the final census. Survival was high between the penultimate and the final censuses; 97% and 85% of individuals alive in March 2013 survived until April 2014 (2005 and 2006 cohorts, respectively). Given that few plants died over the final year, we have not introduced bias into our analyses by coding living plants with the final census date. We also conducted complementary logistic regressions in a generalized linear mixed model framework, analyzing the number of individuals that survived until April 2014 over the number of individuals per family that were initially planted in the study (glmer function of the R package **lme4** ver. 1.1-21, Bates et al. 2015). This logistic regression approach treats all dead individuals identically, whether mortality happened early or late in the experiment.

### Divergent selection

Divergent selection can be detected by a significant interaction between trait and transplant environment in genotypic selection analyses; therefore, we analyzed relative fitness as a function of traits (specific leaf area, leaf size and stomatal density) by transplant habitat with a random effect of genotype.

Across the course of the experiment, mortality was significantly greater in the bottomland than in the upland transplant gardens (Anderson and Geber 2010, and this analysis), such that some families lack trait data for later years because all individuals died. For that reason, we analyzed selection using phenotypic data collected only during the first year of trait measurements (2006 traits for the 2005 cohort, and 2007 traits for the 2006 cohort) for which our trait dataset was the most complete (2005 cohort: N=173 families in upland gardens and N=48 families in bottomland sites; 2006 cohort: N=70 families in upland gardens and N=69 families in bottomland gardens). We modeled viability through April 2014 as a function of these early phenotypic values; thus, we leveraged the full fitness dataset to evaluate selection on traits measured early in the study. Trait values from these years were within the range of trait values expressed in subsequent years (Fig. 1). A restricted dataset focused on the subset of clones for which we had trait data from all sampling time points (2005 cohort: 2006, 2007 and 2012 years; 2006 cohort: 2007 and 2012 years) would lack data on families that died early in the experiment and may have been poorly adapted to upland or bottomland environments (2005 cohort: N=167 families in upland gardens and N=33 families in bottomland gardens; 2006 cohort: N= 61 families in upland gardens and N=36 families in bottomland gardens). Since we evaluated selection across two planting cohorts, we applied a corrected α=0.025 (=0.05/2 non-independent datasets) to assess statistical significance.

**Figure 1:**
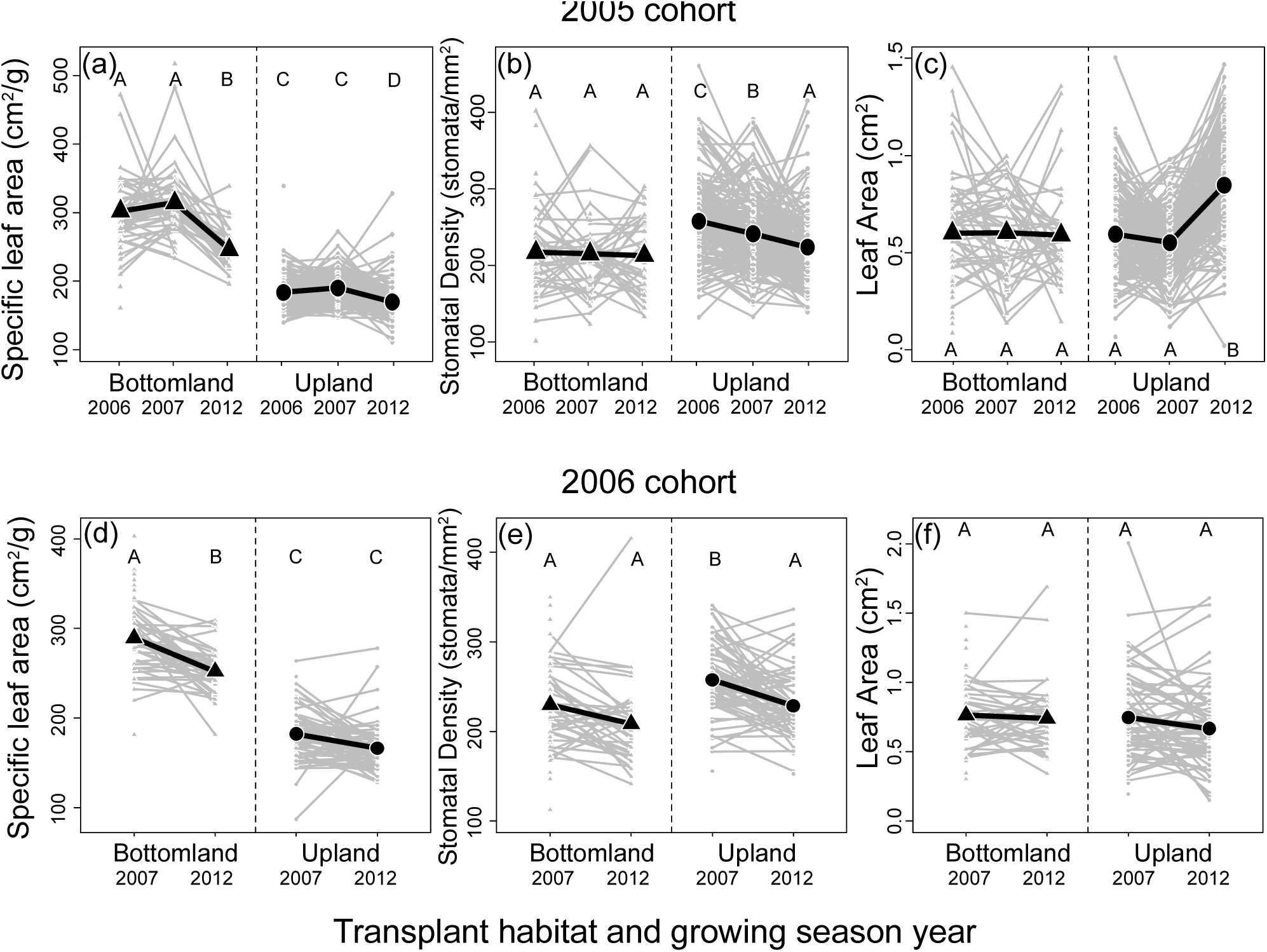
Spatial and temporal variation in (a) stomatal density, (b) leaf area, and (c) specific leaf area for the 2005 cohort measured in 2006, 2007 and 2012, and the same traits (d: SLA; e: stomatal density; f: leaf area) for the 2006 cohort measured in 2007 and 2012. Data points in grey represent LSMEAN trait values for N=183 (2005 cohort) and 79 (2006 cohort) clonal families included in the transplant experiment. The black symbols and lines reflect the overall means across families. We analyzed all traits simultaneously using repeated measures multivariate regression. To achieve model convergence, we standardized traits to a mean of 0 and standard deviation of 1, but we present unstandardized values here. Letters within each panel represent significant differences across habitat types and growing seasons for each trait separately after Tukey’s adjustment for multiple comparison.

In heterogeneous systems subject to demographic source-sink dynamics, analyses of absolute fitness may predominately detect selection in the more frequent or higher quality environment (upland forests) whereas analyses of relative fitness provide more robust information on selection across each of the habitat types (Stanton and Thiede 2005). For that reason, we evaluated soft selection using relative fitness. To calculate relative fitness for longevity, we divided each family’s absolute fitness by the mean fitness expressed by all families in that transplant environment. We tested for nonlinear selection by evaluating quadratic effects of traits and their interactions with environments; we removed any nonsignificant quadratic effects from the final models. For all analyses, we visualized selection using the R package **visreg** vers. 2.6-0 (Breheny and Burchett 2017) by plotting partial residuals from the multiple regressions while holding other explanatory variables at their median value (conditional plots).

### Selection on plasticity

To test whether plasticity is adaptive, we used across-environment multivariate genotypic selection analysis (Van Kleunen and Fischer 2001; Stinchcombe et al. 2004), using the glmer (generalized linear mixed models with gamma distribution) functions of the R package **lme4** (ver. 1.1-21, Bates et al. 2015). As with our analyses of divergent selection, we tested whether selection favored spatial plasticity using the first year of trait data (N=41 families for the 2005 cohort; N=60 families for the 2006 cohort), for which we have a larger sample size than if we restricted the dataset to families with data from all trait sampling points (N=24 families for the 2005 cohort; N=28 families for the 2006 cohort). For these analyses, we also used a corrected α=0.025 (=0.05/2 sets of analyses) to assess statistical significance because we included two cohorts.

In multiple regression analyses, we modeled relative fitness as a function of family-mean trait values (averaged across environments) and plasticity in traits to identify selection on plasticity independent from selection on trait values. For each family, we quantified plasticity via a modified version of the phenotypic plasticity index (PI_LSM_), based on least square mean trait values for each clone in each environment (Valladares et al. 2006). The original PI_LSM_ metric is calculated as the difference between maximum LSMEAN trait values and minimum LSMEAN trait values divided by the maximum LSMEAN trait value [PI_LSM_ = (LSMEAN_maximum_ - LSMEAN_minimum_)/ LSMEAN_maximum_]. This metric quantifies the magnitude, but obscures the directionality of plasticity. In our system, some genotypes express plasticity in the opposite direction from average trait changes across habitat types and seasons, which could be maladaptive. As we aim to test whether plasticity is adaptive, our modified plasticity index incorporates the directionality of plasticity into Valladares et al.’s (2006) framework by quantifying plasticity as: LSMEAN_E, high_ - LSMEAN_E, low_)/ LSMEAN_E, high_, where LSMEAN_E, high_ is the family-mean trait value in the environment with a global average higher mean for that trait, and LSMEAN_E, low_ is the family-mean trait value in the environment with a global lower mean. This formula maintains positive expected plasticity values, but allows for negative values for families that shift their trait values in the opposite direction from the population as a whole. For example, for both cohorts across study years, specific leaf area (SLA) was significantly greater in the bottomland transplant environment than in the upland transplant environment (Fig. 1). Therefore, for this trait: plasticity index= (SLA_bottomland_-SLA_upland_)/ SLA_bottomland_. We used the same configuration for leaf area because it generated an average positive plasticity index for both cohorts. Upland transplants expressed higher stomatal density values than bottomland transplants, so we considered the upland environment to be the minuend in the numerator, and the factor in the denominator in the calculation of plasticity in this trait.

If plasticity confers a fitness advantage in the source (upland) habitat, selection could maintain adaptive plasticity across the landscape despite demographic source-sink dynamics. To test whether selection favors plasticity in the source habitat, we analyzed genotypic fitness (longevity) of clones transplanted into the upland gardens as a function of trait values expressed within upland sites only and spatial plasticity for the first year of trait measurement for both cohorts. We focused on the first year of data to maximize statistical power to test our hypothesis with the largest datasets available. This analysis tests whether the most phenotypically labile plants had greater longevity within upland forests. We ran these generalized linear mixed models using a gamma distribution and log link, and including a random effect for source population, in the glmer function of the R package **lme4** (ver. 1.1-21, Bates et al. 2015).

## Results

### Phenotypic plasticity

We found significant temporal and spatial plasticity for both cohorts (Table S2; Fig. 1). Furthermore, temporal variation was broadly concordant across cohorts. Nevertheless, trait variation was not always congruent with expectations. In the 2005 cohort, analyses confirmed previously reported plasticity in specific leaf area, as well as documenting plasticity in stomatal density and leaf area (Fig. 1a-1c; Table S2: trait **×** transplant habitat **×** season: F_6,513_=38.4, p<0.0001). Specific leaf area (SLA) was significantly lower in upland than bottomland environments across years and SLA values varied with year within habitats (Fig. 1a). Consistent with expectations, stomatal density was higher in upland than bottomland transplant habitats in two of three years; stomatal density varied across years in the upland environment but not in the bottomlands (Fig. 1b). Finally, leaf area was significantly greater in uplands than bottomlands in one year only (Fig. 1c). Significant source habitat by transplant habitat interactions and source habitat by growing season interactions indicated that the magnitude of spatio-temporal plasticity was slightly greater for bottomland than upland genotypes (Table S2), consistent with theoretical predictions of greater plasticity in marginal habitats (Chevin and Lande 2011).

The 2006 cohort also displayed a significant interaction among transplant habitat and growing season for all traits (Fig. 1d-1f; Table S2, trait **×** transplant habitat **×** season: F_3,146_=2.68, p=0.049). Concordant with expectations, experimental individuals expressed greater specific leaf area under the dense canopy of bottomland habitats than in the higher light environment of upland forests (Fig. 1d). Temporal variation in SLA was apparent in the bottomland but not the upland environment (Fig. 1d). Stomatal density was significantly greater in upland forests in one year (2007) than in all other transplant habitat by season combinations (Fig. 1e). However, leaf area did not differ across space or time (Fig. 1f). Finally, for leaf area only, our analyses found a significant effect of source habitat (F_1,208.5_=68.58, p<0.0001), such that upland genotypes had larger leaves than bottomland genotypes (contrast in standardized trait values between bottomland and upland genotypes: -0.43 ± 0.15; Table S2).

### Divergent selection

#### Stomatal density

Divergent selection in the 2005 cohort operated on stomatal density (quadratic trait **×** transplant habitat interaction: χ^2^=9.56, p=0.002, Fig. 2a, Table S3), with stabilizing selection favored an intermediate stomatal density in bottomland forests and no apparent selection within upland habitats. We found no evidence for divergent selection on this trait in the 2006 cohort. Contrary to expectations, logistic regression of survival revealed directional selection for low stomatal density in uplands and high stomatal density in bottomlands for the 2005 cohort (Table S4, Fig. S1a). Directional selection for increased stomatal density in the uplands in the 2006 cohort accorded with predictions (Table S4, Fig. S1b).

**Figure 2:**
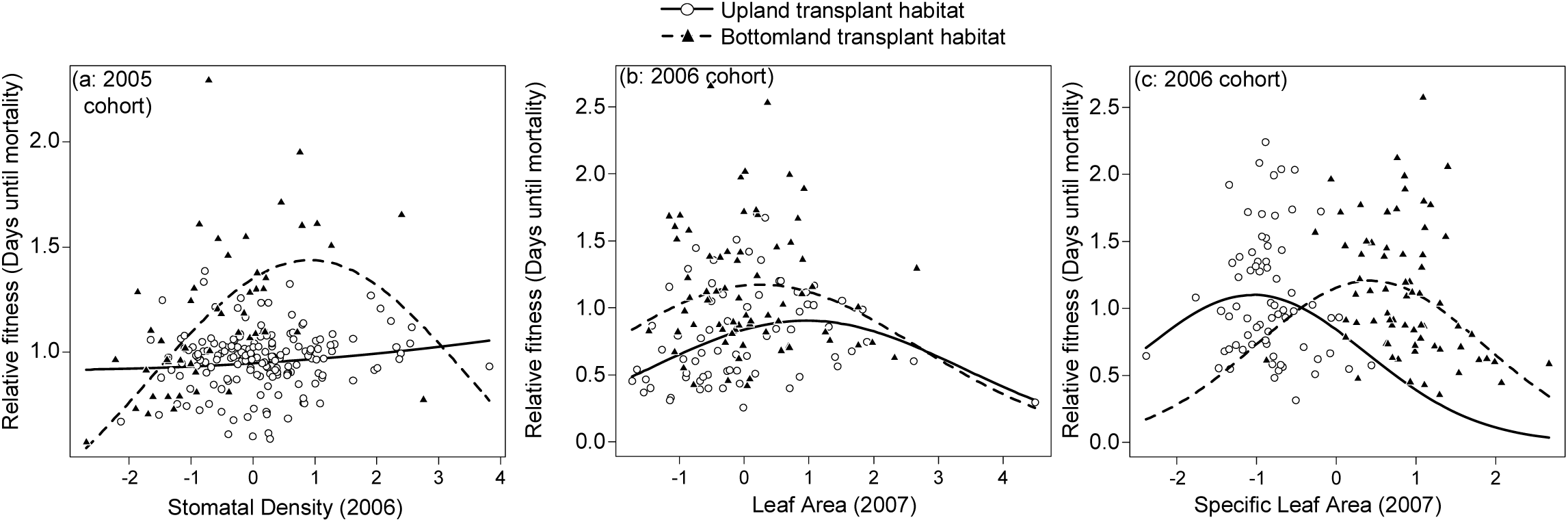
Longevity selection on foliar traits in the 2005 and 2006 cohorts. Gamma regression analyses revealed selection operating on stomatal density (a) during the first year of trait measurement for the 2005 cohort. Stabilizing selection favored intermediate leaf are for (b) the 2006 cohort for both habitats. Finally, (c) divergent selection on specific leaf area in the 2006 cohort was consistent with the direction of plasticity. Bottomland data and predictedregression lines are displayed with closed triangles and dashed lines; upland data and regression lines are shown with open circlesand solid lines. Traits were standardized to a mean of 0 and standard deviation of 1. Panels show partial residuals from multiple regressions while holding other traits at their median value.

#### Leaf area

For the 2006 cohort: stabilizing longevity selection favored intermediate leaf sizes in the first year of measurement in both habitats (quadratic trait: χ^2^=12.98, p=0.00031, Fig. 2b, Table S3). Logistic regression revealed viability selection for increased leaf size in upland forests in both cohorts, and smaller leaf size in the bottomland forests in the 2006 cohort (Fig. S1c,d, Table S4).

#### Specific leaf area (SLA)

Concordant with expectations, selection favored lower specific leaf area in the upland forests and higher specific leaf area in the bottomland forests in the 2006 cohort (SLA **×** transplant habitat interaction: χ^2^=5.15, p=0.023; quadratic trait: χ^2^ =10.14, p=0.0015; Fig. 2c, Table S3). We did not detect longevity selection on SLA for the 2005 cohort (Table S3). However, for both cohorts, divergent viability selection assessed via logistic regression favored reduced SLA in uplands and increased SLA in bottomland forests (Fig. S1e, f, Tables S4), consistent with the direction of trait plasticity (Fig. 1a and d).

### Selection on plasticity

#### Stomatal density

Nonlinear selection favored intermediate plasticity in stomatal density for the 2006 cohort (plasticity optimum: 0.10; quadratic effect: χ^2^=8.3, p=0.0039; Fig. 3a, Table S5). We did not find viability selection on plasticity in stomatal density (Table S6).

**Figure 3:**
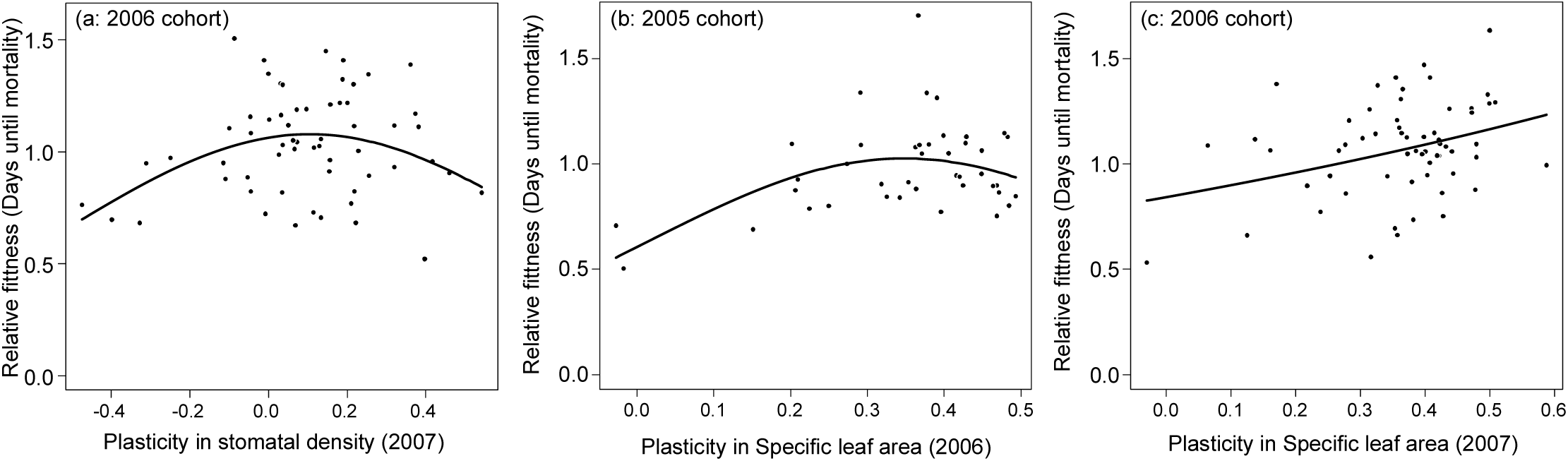
Longevity selection acted on plasticity in foliar traits in both cohorts. Stabilizing selection favored intermediate plasticity in (a) stomatal density for the 2006 cohort, and (b) specific leaf area in the 2005 cohort. Directional selection favored (c) increased plasticity in specific leaf area for the 2005 cohort. These multivariate genotypic selection analyses evaluated fitness as a function of mean trait values and plasticities in all three traits, with separate models for each planting cohort. Panels show partial residuals from multiple regressions while holding other traits at their median value.

#### Leaf area

We found no evidence for selection on leaf area plasticity in either cohort (Tables S5 and S6).

#### Specific leaf area (SLA)

For the 2005 cohort, stabilizing selection operated on specific leaf area (plasticity optimum: 0.35; quadratic effect: χ^2^ =6.35, p=0.012; Fig. 3b, Table S5). For the 2006 cohort, directional selection favored increased plasticity in specific leaf area (χ^2^=6.34, p=0.012; Fig. 3c, Table S5).

#### Selection on plasticity within the source environment

Across both cohorts, longevity selection favored greater plasticity in stomatal density and specific leaf area (Table S7, Fig. 4). For the 2005 cohort, fitness increased with plasticity in stomatal density (χ^2^=8.65, p=0.0033; Fig. 4a). A negative quadratic curve suggested stabilizing selection on plasticity in stomatal density in the 2006 cohort, favoring intermediate values (plasticity optimum: 0.13, quadratic term: χ^2^=5.34, p=0.021; Fig. 4b). We detected a trend that could reflect nonlinear selection for greater plasticity in specific leaf area in the 2005 cohort (linear term: χ^2^=5.36, p=0.021; quadratic term: χ^2^=3.55, p=0.06; Fig. 4c). Directional selection within the uplands favored increased plasticity in specific leaf area in the 2006 cohort (χ^2^=7.38, p=0.0066; Fig. 4d).

**Figure 4:**
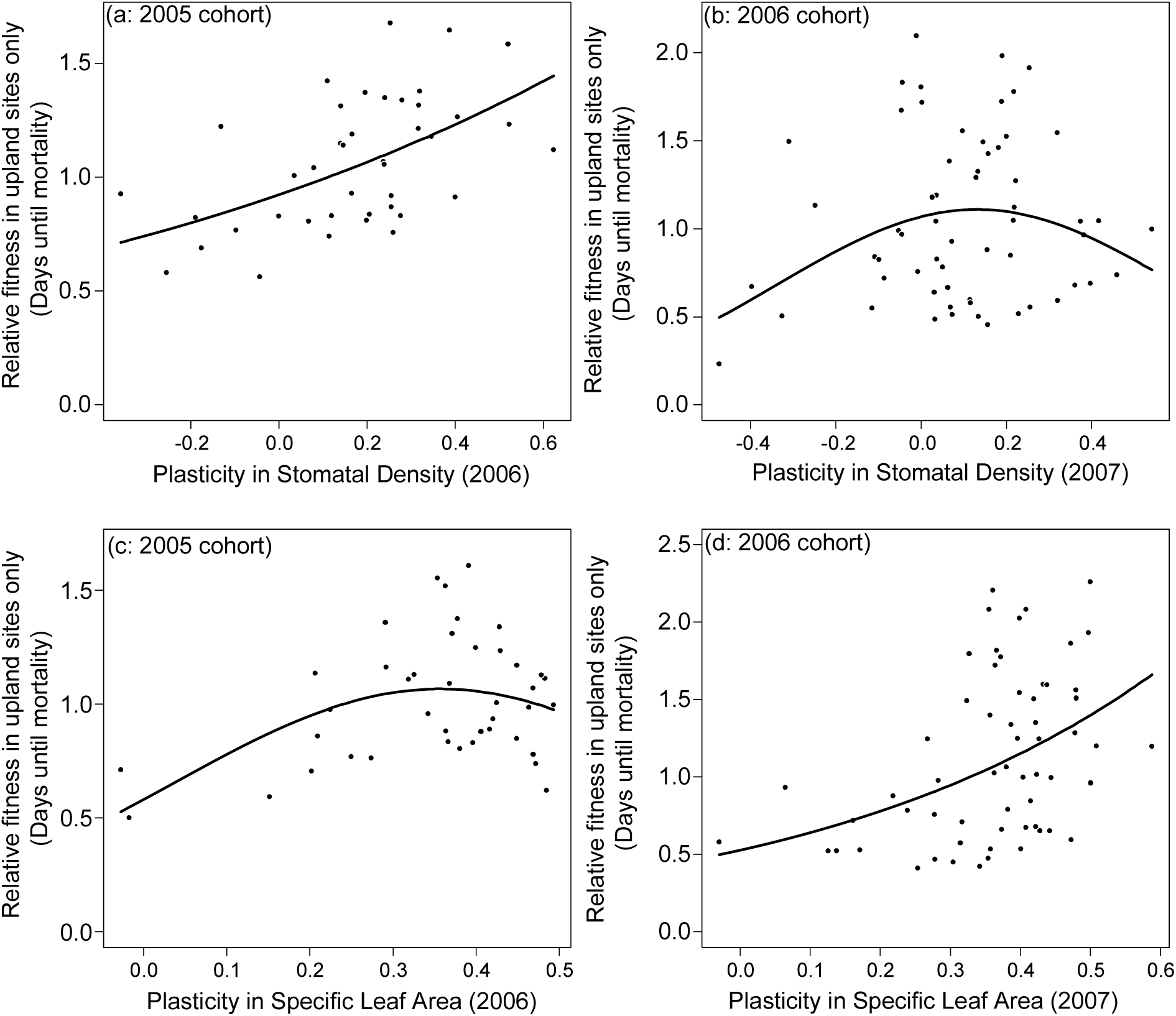
Selection favored plasticity in stomatal density (a: 2005 cohort; b: 2006 cohort) and specific leaf area (c: 2005 cohort; d: 2006 cohort) within upland forests. Panels show partial residuals from multiple regressions while holding other traits at their median value. Data points represent relative fitness based on genotype mean longevity as a function of spatial plasticity in traits measured in the first year for each cohort. Panels show partial residuals from multiple regressions while holding other traits at their median value.

## Discussion

When individuals express higher fitness in one habitat relative to another, selection is expected to favor trait values advantageous in the higher-quality or larger habitat at the expense of adaptations to the lower-quality habitat (Holt and Gaines 1992; Stanton and Thiede 2005; Kawecki 2008). Similarly, if populations inhabiting a marginal environment produce fewer propagules than those from a higher quality habitat, evolution might not favor a plastic response to spatial heterogeneity (Holt and Gaines 1992; Sultan and Spencer 2002). Nevertheless, in our study, adaptive plasticity in morphological traits conferred a fitness advantage for *Vaccinium elliottii* across environments. In addition, selection favored adaptive plasticity in stomatal density and specific leaf area within upland forests (Fig. 4) where *V*. *elliotti* individuals have greater survival in our reciprocal transplant experiment, and natural populations are more fecund and abundant (Anderson and Geber 2010). We hypothesize that plasticity could be advantageous within upland forests because of microenvironmental spatial heterogeneity and temporal variation in conditions. Selection within upland forests could maintain adaptive morphological plasticity across populations in both habitat types. Further, we hypothesize that this plasticity - driven by selection in the uplands - could enhance survival within the lower-quality bottomland habitat, as is proposed in theoretical models (Chevin and Lande 2011). Our analyses support the hypothesis that selection can favor adaptive plasticity in a spatially and temporally heterogeneous landscape, despite demographic source-sink dynamics. Indeed, this adaptive plasticity could reduce maladaptation and enhance fitness in the marginal environment (Chevin and Lande 2011).

### Divergent selection

Based on trait variation across habitat types in this system (Fig. 1) in concert with clinal variation in other systems (Steinger et al. 2003; Wright et al. 2004; Carlson et al. 2015; Maire et al. 2015; Ramírez-Valiente et al. 2018), we expected selection to favor increased specific leaf area and leaf area, and reduced stomatal density in bottomland relative to upland forests. Indeed, concordant with predictions, we found divergent longevity (and viability) selection on specific leaf area. In contrast, selection on stomatal density did not follow expectations. Rather, in the 2006 cohort, stabilizing selection favored intermediate stomatal density in bottomlands, but we found no evidence for selection in upland habitats.

We expected to find smaller leaves in upland than bottomland forests because leaf lamina area often declines with increasing aridity and light levels (Valladares et al. 2000; Carlson et al. 2015; Ramírez-Valiente et al. 2018). Instead, we found the opposite pattern: leaves were similar in size across habitats in two years (2006 and 2007) and larger in upland than bottomland environments in another year (2012). Our analysis detected similar patterns of stabilizing selection on leaf size in both habitats. Not surprisingly, we found no evidence that selection favors plasticity in leaf area, which was the least plastic trait in the study and which is not subject to divergent selection across habitat types. Trait expression and selection on leaf area could be driven by factors other than aridity or understory light levels in this system. We note that our models incorporate indirect selection on focal traits mediated by unmeasured traits.

### Selection for adaptive plasticity

Spatial and temporal heterogeneity in environmental conditions can promote the evolution of adaptive plasticity when individuals experience multiple environmental conditions across their lifetimes or when the progeny disperse into non-parental habitat types (Baythavong and Stanton 2010; Baythavong 2011). Models suggest that even low levels of gene flow can favor the evolution of phenotypic plasticity (Sultan and Spencer 2002). In our system, asymmetric gene flow occurs predominately from upland to bottomland populations, yet rare gene flow in the reverse direction also connects populations (Anderson and Geber 2010). Given the high rates of gene flow across habitat types (Anderson and Geber 2010), *V*. *elliottii* seeds likely often germinate and establish in different environments than their maternal and paternal parents. In addition, water stress can vary inter- and intra-annually in both bottomland and upland habitats. Thus, in both habitat types, established individuals experience multiple years of variable environmental conditions prior to reproduction, which could strongly favor the evolution of adaptive plasticity in functional traits.

Evolutionary studies that have explicitly tested the adaptive significance of plasticity in plants focus primarily on herbaceous systems (Dudley and Schmitt 1996; Scheiner and Callahan 1999; Schmitt et al. 1999; Donohue et al. 2000; Steinger et al. 2003; Bell and Galloway 2007; Galloway and Etterson 2007; Baythavong 2011; Zhang et al. 2013; Wagner and Mitchell-Olds 2018), even though woody plants represent ∼45-48% of plant species globally (FitzJohn et al. 2014). Woody plant species typically have reduced population genetic structure (lower F_ST_) relative to annual or perennial herbaceous species, indicating greater rates of gene flow (Duminil et al. 2009). For those reasons, plasticity could be particularly adaptive for woody species, because they may be more likely than herbaceous species to experience temporal and spatial variation in environmental conditions. Even though our study of post-establishment survival did not capture the full extent of selection operating across the duration of the life cycle, we found that adaptive plasticity in stomatal density and specific leaf area confers a viability and longevity advantage across multiple years.

### Stabilizing selection on plasticity

An additional pattern emerged in our study: Stabilizing selection favored intermediate plasticity in stomatal density and specific leaf area. We propose that families with low levels of trait plasticity might not express appropriate phenotypes in response to spatial or temporal variation in environmental conditions. Similarly, families with very high levels of plasticity could be too labile, perhaps shifting phenotypes too readily or expressing exaggerated trait values. Thus, we might expect fitness to be maximized at some intermediate trait plasticity, just as stabilizing selection can favor intermediate trait expression in multivariate trait space (e.g., Brooks et al. 2005; Wadgymar et al. 2017; Taylor et al. 2018). Logistical constraints often preclude the sample sizes necessary to gain sufficient statistical power for these analyses and the duration of the field studies from which to estimate plasticity and fitness. We suspect that as statistical tools become more powerful, researchers will uncover more examples of nonlinear selection on plasticity.

### Selection varies across cohorts

The magnitude and direction of selection can change through time. In several instances, the degree of selection differed across cohorts. In our experiment, the 2005 cohort experienced a large-scale flood event in the bottomland gardens during planting, which contrasted with the average conditions experienced by the 2006 cohort during and shortly after planting. Extreme events, such as the flood of 2005, can impose strong viability selection, which can restrict the number of individuals that survive these events and influence the distribution of trait and fitness values of the survivors. Our results indicate that conditions during initial establishment can set the stage for trait expression and selection later in life history.

### Conclusions

Our analyses suggest that spatial and temporal variation in environmental conditions favors phenotypic plasticity. Demographic source-sink dynamics pose challenges for conservation in contemporary landscapes, as small disconnected habitat patches can be associated with poor performance (Furrer and Pasinelli 2016) and habitat fragmentation could shift patches from sources to sinks. Understanding eco-evolutionary dynamics in source-sink systems could lead to better conservation outcomes. Asymmetrical gene flow from upland into bottomland forests likely constrains other adaptations to flooding such as adventitious roots, porous root tissue, and the formation of enlarged lenticels, which are present in other species of *Vaccinium* (Anderson and Geber 2010). Nevertheless, selection operating across this heterogeneous landscape can favor plasticity in key functional traits, which could enhance fitness and population persistence within the marginal bottomland habitat type (Chevin and Lande 2011). We suggest that phenotypic plasticity is likely to be advantageous in other systems when individuals encounter multiple environments over their lifetimes and progeny disperse to non-parental environments.

## Supplemental materials

**Table S1:** Trait correlations in upland and bottomland transplant habitats for the 2005 cohort. We find relatively low levels of correlations across traits in this experiment. Below, we include the Pearson correlation coefficients and (uncorrected) p-values for trait correlations in both transplant habitat types for the 2005 cohort. The three foliar traits are sometimes correlated with each other, but these correlations vary substantially. We have not applied any corrections for multiple testing in these tables. Trait correlations are similar for the 2006 cohort and can be calculated from the Dryad datafile associated with this manuscript.

**Upland transplant habitat (2005 cohort):**
|  |  | Stomatal<br>Density,<br>2006 | Specific<br>Leaf Area,<br>2006 | Leaf<br>Area,<br>2006 | Stomatal<br>Density,<br>2007 | Specific<br>Leaf<br>Area,<br>2007 | Leaf<br>area,<br>2007 | Stomatal<br>Density,<br>2012 | Specific<br>Leaf<br>Area,<br>2012 | Leaf<br>area,<br>2012 |
| --- | --- | --- | --- | --- | --- | --- | --- | --- | --- | --- |
| Stomatal Density,<br>2006 | r | 1 |  |  |  |  |  |  |  |  |
|  | p-value |  |  |  |  |  |  |  |  |  |
| Specific Leaf Area,<br>2006 | r | -0.05 | 1 |  |  |  |  |  |  |  |
|  | p-value | 0.5495 |  |  |  |  |  |  |  |  |
| Leaf Area, 2006 | r | -0.11 | -0.22 | 1 |  |  |  |  |  |  |
|  | p-value | 0.1689 | **0.0039** |  |  |  |  |  |  |  |
| Stomatal Density,<br>2007 | r | 0.5 | -0.01 | -0.02 | 1 |  |  |  |  |  |
|  | p-value | **<0.0001** | 0.866 | 0.8401 |  |  |  |  |  |  |
| Specific Leaf Area,<br>2007 | r | -0.08 | 0.47 | -0.12 | -0.05 | 1 |  |  |  |  |
|  | p-value | 0.3089 | 0 | 0.1153 | 0.5115 |  |  |  |  |  |
| Leaf area, 2007 | r | 0.03 | 0.02 | 0.33 | -0.07 | 0.13 | 1 |  |  |  |
|  | p-value | 0.7283 | 0.7668 | **<0.0001** | 0.3278 | 0.0947 |  |  |  |  |
| Stomatal Density,<br>2012 | r | 0.26 | 0.04 | 0 | 0.28 | 0 | 0.05 | 1 |  |  |
|  | p-value | **0.0006** | 0.5834 | 0.9685 | **0.0003** | 0.9703 | 0.512 |  |  |  |
| Specific Leaf Area,<br>2012 | r | 0.05 | 0.31 | -0.1 | 0.01 | 0.28 | -0.05 | 0.18 | 1 |  |
|  | p-value | 0.5092 | **<0.0001** | 0.1838 | 0.939 | **0.0002** | 0.5425 | **0.0203** |  |  |
| Leaf area, 2012 | r | 0 | 0 | 0.32 | -0.05 | 0 | 0.39 | -0.08 | -0.22 | 1 |
|  | p-value | 0.9567 | 0.9995 | **<0.0001** | 0.5046 | 0.9834 | **<0.0001** | 0.2723 | **0.0043** |  |

**Bottomland transplant habitat (2005 cohort):**
|  |  | Stomatal<br>Density,<br>2006 | Specific<br>Leaf Area,<br>2006 | Leaf<br>Area,<br>2006 | Stomatal<br>Density,<br>2007 | Specific<br>Leaf<br>Area,<br>2007 | Leaf<br>area,<br>2007 | Stomatal<br>Density,<br>2012 | Specific<br>Leaf<br>Area,<br>2012 | Leaf<br>area,<br>2012 |
| --- | --- | --- | --- | --- | --- | --- | --- | --- | --- | --- |
| Stomatal<br>Density, 2006 | r | 1 |  |  |  |  |  |  |  |  |
|  | p-value |  |  |  |  |  |  |  |  |  |
| Specific Leaf<br>Area, 2006 | r | 0.12 | 1 |  |  |  |  |  |  |  |
|  | p-value | 0.4229 |  |  |  |  |  |  |  |  |
| Leaf Area, 2006 | r | -0.04 | -0.23 | 1 |  |  |  |  |  |  |
|  | p-value | 0.7674 | 0.0701 |  |  |  |  |  |  |  |
| Stomatal<br>Density, 2007 | r | 0.41 | 0 | -0.24 | 1 |  |  |  |  |  |
|  | p-value | **0.0089** | 0.9886 | 0.1341 |  |  |  |  |  |  |
| Specific Leaf<br>Area, 2007 | r | -0.34 | 0 | -0.07 | 0.17 | 1 |  |  |  |  |
|  | p-value | **0.0237** | 0.9957 | 0.627 | 0.2699 |  |  |  |  |  |
| Leaf area, 2007 | r | 0.09 | 0.06 | 0.33 | -0.1 | -0.27 | 1 |  |  |  |
|  | p-value | 0.5693 | 0.6604 | 0.0209 | 0.5094 | 0.0512 |  |  |  |  |
| Stomatal<br>Density, 2012 | r | 0.45 | -0.15 | 0.03 | 0.25 | -0.05 | -0.22 | 1 |  |  |
|  | p-value | **0.0083** | 0.3936 | 0.851 | 0.16 | 0.754 | 0.1984 |  |  |  |
| Specific Leaf<br>Area, 2012 | r | -0.05 | 0.29 | 0 | -0.17 | 0.21 | -0.11 | 0.03 | 1 |  |
|  | p-value | 0.7828 | 0.0757 | 0.9858 | 0.3077 | 0.2015 | 0.5038 | 0.8764 |  |  |
| Leaf area, 2012 | r | -0.04 | 0.06 | 0.19 | 0 | -0.02 | 0.05 | -0.44 | 0.05 | 1 |
|  | p-value | 0.8142 | 0.7042 | 0.266 | 0.9798 | 0.8879 | 0.7651 | **0.0089** | 0.7562 |  |

**Table S2:** Repeated measures multivariate analyses of stomatal anatomy, specific leaf area, and leaf size from the 2005 and 2006 cohorts demonstrates phenotypic plasticity across time and space (habitat type). These models simultaneously evaluate all three traits and their interactions with growing season, transplant habitat, and source habitat (Proc Mixed, SAS ver. 9.4). Significant interactions between phenotype and other explanatory variables indicate that effects of habitat, season, life history and their interactions differ by trait. We used slice statements in SAS to examine plasticity separately for each trait. We assessed significance of the random effect of genotype and genotype by habitat via likelihood ratio tests by comparison of models with and without these effects (^2^, degrees of freedom = 1). Our analysis of the 2005 cohort uncovered two unexpected interactions with source habitat. The interaction between transplant habitat and source habitat revealed that bottomland genotypes expressed greater spatial plasticity than upland genotypes for all three foliar traits. Here, we present plasticity as tthe contrast in standardized trait values between upland and bottomland transplant sites : stomatal density (F_3,501.3_=7.89, p<0.0001; plasticity of bottomland genotypes : -0.60 ± 0.16, t_472.8_=-3.83, Tukey’s adjusted p=0.0009; plasticity of upland genotypes: -0.44 ± 0.14, t_559.5_=-3.00, p=0.015); specific leaf area (F_3,235.6_=198.5, p<0.0001, plasticity of bottomland genotypes: 1.96 ± 0.1, t_237.1_=18.6, p<0.0001; than upland genotypes plasticity: 1.65 ± 0.1, t_216.8_=15.8, p<0.0001); and leaf area (F_3,513.3_=5.39, p=0.0012; plasticity of bottomland genotypes: -0.53 ± 0.14; t_490_= -3.81, p=0.0009; plasticity of upland genotypes: -0.078 ± 0.14; t_458_=-0.57, p=0.94). The interaction between source habitat and growing season revealed that bottomland genotypes expressed greater temporal plasticity than upland genotypes for two of the three foliar traits. For example, for stomatal density (F_5,490_=3.81, p=0.0021), plasticity between the year with the greatest stomatal density (2006) and the year with the lowest average stomatal density (2012) was greater for bottomland genotypes (0.53 ± 0.15; t_375.6_=3.58, p=0.0051) than upland genotypes (0.30 ± 0.14; t_361.7_=2.24, p=0. 22). Similarly, bottomland genotypes had greater temporal plasticity than upland genotypes in leaf area (F_5,516.5_=10.1, p<0.0001; plasticity of bottomland genotypes from 2006 to 2012: -0.43 ± 0.13; t_406.9_=-3.34, p=0. 012; plasticity of upland genotypes: -0.31 ± 0.12; t_415.3_=-2.51, p=0.12). Finally, the source habitat by season interaction for specific leaf area (F_5,436_=48.18, p<0.0001) was not as straightforward. When we evaluated years separately, we found no difference in traits values between source habitats in 2006 (t_394.5_=0, p=1), 2007 (t_445.8_=0.91, p=0.94), or 2012 (t_566.1_=1.1, p=0.88). Therefore, this interaction may have arisen through only very slight shifts in the rankings of genotypes across seasons. For example, the shift in trait values between the year with the greatest SLA (2006) and the year with the lowest average SLA (2012) was slightly lower for bottomland genotypes (contrast in standardized trait values between 2006 and 2012: 0.56 ± 0.07; t_267.4_=7.6, p<0.0001) than upland genotypes (contrast in standardized trait values between 2006 and 2012: 0.67 ± 0.07; t_261.4_=9.42, p=<0.0001).

| Effect | 2005 cohort |  | 2006 cohort |  |
| --- | --- | --- | --- | --- |
|  | F-value | p-value | F-value | p-value |
| Phenotype (P) | $F_{3,421}=49.4$ | **<0.0001** | $F_{3,163}=0.53$ | 0.66 |
| $P \times \text{Season}$ | $F_{6,513}=43.3$ | **<0.0001** | $F_{3,146}=17.22$ | **<0.0001** |
| $P \times \text{Transplant habitat}$ | $F_{3,376}=209.7$ | **<0.0001** | $F_{3,111}=62.7$ | **<0.0001** |
| $P \times \text{Source habitat}$ | $F_{3,421}=0.43$ | 0.73 | $F_{3,163}=3.33$ | **0.0211** |
| $P \times \text{Transplant habitat} \times \text{Season}$ | $F_{6,513}=28.4$ | **<0.0001** | $F_{3,146}=2.68$ | **0.049** |
| $P \times \text{Source habitat} \times \text{Season}$ | $F_{6,513}=2.48$ | **0.023** | $F_{3,146}=1.51$ | 0.21 |
| $P \times \text{Transplant habitat} \times \text{Source habitat}$ | $F_{3,376}=3.3$ | **0.021** | $F_{3,111}=0.92$ | 0.43 |
| $P \times \text{Transplant habitat} \times \text{Source habitat} \times \text{Season}$ | $F_{6,513}=1.23$ | 0.29 | $F_{3,146}=0.41$ | 0.75 |

| | $\chi^2$ | p-value | $\chi^2$ | p-value |
| --- | --- | --- | --- | --- |
| Genotype | 8.3 | **0.004** | 12.8 | **0.00035** |
| Genotype × Transplant habitat | 18.2 | **<0.0001** | 0 | 1 |

**Table S3:** Results of longevity selection (relative days until mortality) models of on foliar phenotypes for the 2005 and 2006 cohorts. We assessed significance of the random effect of genotype via likelihood ratio tests (χ^2^, degrees of freedom = 1). We used a corrected α=0.025 (=0.05/2 sets of analyses) to assess statistical significance. For significant traits, we present partial regression coefficients ± standard errors to evaluate the magnitude of selection. It is important to consider that we present unexponentiated coefficients from these Gamma regressions, and undoubled quadratic coefficients. To estimate quadratic selection gradients (γ), partial regression coefficients and standard errors must be doubled (Stinchcombe et al. 2008).

|  | 2005 cohort |  |  |  | 2006 cohort |  |  |  |
| --- | --- | --- | --- | --- | --- | --- | --- | --- |
| | Partial regression coefficients ± S.E. | $\chi^2$ | Df | p-value | Partial regression coefficients ± S.E. | $\chi^2$ | Df | p-value |
| Transplant habitat | NA | 7.795 | 1 | **<0.0001** | NA | 3.703 | 1 | **<0.0001** |
| Specific leaf area | NA | 0.007 | 1 | 0.932 | See transplant habitat by Specific Leaf Area coefficients for each habitat | 7.80 | 1 | **0.005** |
| Stomatal Density | NA | 0.347 | 1 | 0.556 | NA | 0.12 | 1 | 0.73 |
| Leaf area | NA | 0.579 | 1 | 0.447 | 0.10 ± 0.05 | 7.78 | 1 | **0.005** |
| Transplant habitat × Specific Leaf Area | NA | 1.276 | 1 | 0.259 | Upland:<br>-0.52 ± 0.18<br>Bottomland:<br>0.21 ± 0.17 | 5.15 | 1 | **0.023** |
| Transplant habitat × Stomatal Density | Upland:<br>0.019 ± 0.032<br>Bottomland:<br>0.14 ± 0.034 | 9.027 | 1 | **0.0027** | NA | 1.82 | 1 | 0.18 |
| Transplant habitat × Leaf Area | NA | 2.839 | 1 | 0.092 | NA | 2.29 | 1 | 0.13 |
| Stomatal density, quadratic effect | NA | 0.026 | 1 | 0.873 | NA | NA | NA | NA |
| Transplant habitat × Stomatal Density (quadratic) | Upland:<br>0.0026 ± 0.016<br>Bottomland:<br>-0.074 ± 0.022 | 9.564 | 1 | **0.0020** | NA | NA | NA | NA |
| Leaf area, quadratic effect | NA | NA | 1 | NA | -0.085 ± 0.023 | 12.984 | 1 | **0.00031** |
| Specific Leaf area, quadratic effect | NA | NA | 1 | NA | -0.25 ± 0.078 | 10.138 | 1 | **0.0015** |
| Genotype | NA | 88.9 | 1 | **<0.0001** | NA | 59.5 | 1 | **<0.0001** |

**Table S4:** Results of logistic regression models of viability (# individuals alive in April 2014/# individuals planted per family) for the 2005 and 2006 cohorts. We assessed significance of the random effect of genotype via likelihood ratio tests (χ^2^, degrees of freedom = 1). We used an adjusted α=0.025 (=0.05/2 traits) to correct for multiple testing.

|  | 2005 cohort |  |  | 2006 cohort |  |  |
| --- | --- | --- | --- | --- | --- | --- |
| | $\chi^2$ | Df | p-value | $\chi^2$ | Df | p-value |
| Transplant habitat | 15.8 | 1 | **<0.0001** | 0.98 | 1 | 0.32 |
| Specific leaf area | 2.06 | 1 | 0.15 | 6.93 | 1 | **0.0085** |
| Stomatal Density | 2.80 | 1 | 0.09 | 7.07 | 1 | **0.0079** |
| Leaf area | 9.61 | 1 | **0.0019** | 15.66 | 1 | **<0.0001** |
| Transplant habitat × Specific Leaf Area | 10.71 | 1 | **0.0011** | 3.35 | 1 | 0.067 |
| Transplant habitat × Stomatal Density | 7.39 | 1 | **0.0066** | 4.33 | 1 | 0.037 |
| Transplant habitat × Leaf Area | 2.4 | 1 | 0.12 | 7.13 | 1 | **0.0076** |
| Leaf area, quadratic effect | NA | NA | NA | 4.26 | 1 | 0.039 |
| Specific leaf area, quadratic effect | NA | NA | NA | 4.57 | 1 | 0.033 |
| Genotype | 0.10 | 1 | 0.75 | 3.72 | 1 | 0.054 |

**Table S5:** Selection on plasticity for the 2005 and 2006 cohorts. Analyses evaluated selection in models that included trait values averaged across environments and plasticity in those traits. We incorporated quadratic effects of plasticity terms if preliminary models indicated nonlinear selection. We modeled population of origin as a random effect, assessing significance via likelihood ratio tests (χ^2^, degrees of freedom = 1). We used an adjusted α=0.025 (=0.05/2 traits) to correct for multiple testing across two cohorts. For significant traits, we present partial regression coefficients ± standard errors to evaluate the magnitude of selection. It is important to consider that we present unexponentiated coefficients from these Gamma regressions, and undoubled quadratic coefficients. To estimate quadratic selection gradients (γ), partial regression coefficients and standard errors must be doubled (Stinchcombe et al. 2008).

|  | 2005 cohort |  |  |  | 2006 cohort |  |  |  |
| --- | --- | --- | --- | --- | --- | --- | --- | --- |
| | Partial regression coefficients ± S.E. | $\chi^2$ | Df | p-value | Partial regression coefficients ± S.E. | $\chi^2$ | Df | p-value |
| Mean Specific leaf area across environments | NA | 1.011 | 1 | 0.315 | NA | 1.36 | 1 | 0.2 |
| Mean Stomatal Density, across environments | NA | 0.182 | 1 | 0.67 | NA | 1.17 | 1 | 0.2 |
| Mean Leaf area, across environments | NA | 0.128 | 1 | 0.721 | NA | 0.011 | 1 | 0.9 |
| Plasticity in Specific Leaf Area | $3.03 \pm 1.12$ | 7.339 | 1 | **0.0067** | $0.65 \pm 0.26$ | 6.38 | 1 | **0.01** |
| Plasticity in Stomatal Density | NA | 0.247 | 1 | 0.619 | $0.27 \pm 0.15$ | 3.21 | 1 | 0.07 |
| Plasticity in Leaf Area | NA | 2.542 | 1 | 0.111 | NA | 2.57 | 1 | 0.1 |
| Plasticity in Specific Leaf Area, quadratic effect | $-4.26 \pm 1.8$ | 6.347 | 1 | **0.012** | NA | NA | NA | N. |
| Plasticity in Stomatal Density, quadratic effect | NA | NA | NA | NA | $-1.30 \pm 0.45$ | 8.31 | 1 | **0.003** |
| Source population | NA | 1.89 | 1 | 0.17 | NA | 0 | 1 |  |

**Table S6:** Selection on plasticity for the 2005 and 2006 cohorts via viability (# individuals alive in April 2014/# individuals planted per family). Genotypic selection analyses evaluated selection in models that included mean trait values and plasticity in those traits. We incorporated quadratic effects of plasticity terms if preliminary models indicated nonlinear selection. We used an adjusted α=0.025 (=0.05/2 traits) to correct for multiple testing across two cohorts.

|  | 2005 cohort |  |  | 2006 cohort |  |  |
| --- | --- | --- | --- | --- | --- | --- |
| | $\chi^2$ | Df | p-value | $\chi^2$ | Df | p-value |
| Specific leaf area | 1.28 | 1 | 0.26 | 0.44 | 1 | 0.51 |
| Stomatal Density | 0.079 | 1 | 0.78 | 0.026 | 1 | 0.87 |
| Leaf area | 0.31 | 1 | 0.58 | 0.5 | 1 | 0.48 |
| Plasticity in Specific Leaf area | 1.33 | 1 | 0.25 | 0.15 | 1 | 0.7 |
| Plasticity in Stomatal Density | 0.003 | 1 | 0.96 | 1.97 | 1 | 0.16 |
| Plasticity in Leaf Area | 0.012 | 1 | 0.91 | 1.15 | 1 | 0.28 |
| Source population | 0 | 1 | 1 | 0 | 1 | 1 |

**Table S7:** Selection on plasticity for the 2005 and 2006 cohorts within upland forests only. Analyses evaluated longevity within the uplands as a function of trait values averaged within the upland and spatial plasticity in those traits. We incorporated quadratic effects of plasticity terms if preliminary models indicated nonlinear selection. We modeled population of origin as a random effect, assessing significance via likelihood ratio tests (χ^2^, degrees of freedom = 1). We used an adjusted α=0.025 (=0.05/2 traits) to correct for multiple testing across two cohorts. For significant traits, we present partial regression coefficients ± standard errors to evaluate the magnitude of selection. It is important to consider that we present unexponentiated coefficients from these Gamma regressions, and undoubled quadratic coefficients. To estimate quadratic selection gradients (γ), partial regression coefficients and standard errors must be doubled (Stinchcombe et al. 2008).

|  | 2005 cohort |  |  |  | 2006 cohort |  |  |  |
| --- | --- | --- | --- | --- | --- | --- | --- | --- |
| | Partial regression coefficients ± S.E. | $\chi^2$ | Df | p-value | Partial regression coefficients ± S.E. | $\chi^2$ | Df | p-value |
| Specific leaf area, upland average | NA | 2.5 | 1 | 0.114 | NA | 0.84 | 1 | 0.3 |
| Stomatal Density, upland average | NA | 1.71 | 1 | 0.19 | NA | 0.58 | 1 | 0.4 |
| Leaf area, upland average | NA | 0.37 | 1 | 0.544 | NA | 0.26 | 1 | 0.6 |
| Plasticity in Specific Leaf area | $3.4 \pm 1.5$ | 5.36 | 1 | **0.0206** | $1.95 \pm 0.72$ | 7.38 | 1 | **0.006** |
| Plasticity in Stomatal Density | $0.72 \pm 0.24$ | 8.65 | 1 | **0.0033** | $0.58 \pm 0.34$ | 2.93 | 1 | 0.087 |
| Plasticity in Leaf Area | NA | 2.61 | 1 | 0.106 | NA | 0.02 | 1 | 0. |
| Plasticity in Specific Leaf area, quadratic | $-4.8 \pm 2.5$ | 3.55 | 1 | 0.0596 | NA | NA | NA | N. |
| Plasticity in Stomatal density, quadratic | NA | NA | NA | NA | $-2.2 \pm 0.95$ | 5.34 | 1 | **0.020** |
| Population of origin | NA | 2.95 | 1 | 0.086 | NA | 0 | 1 |  |

**Figure S1:**
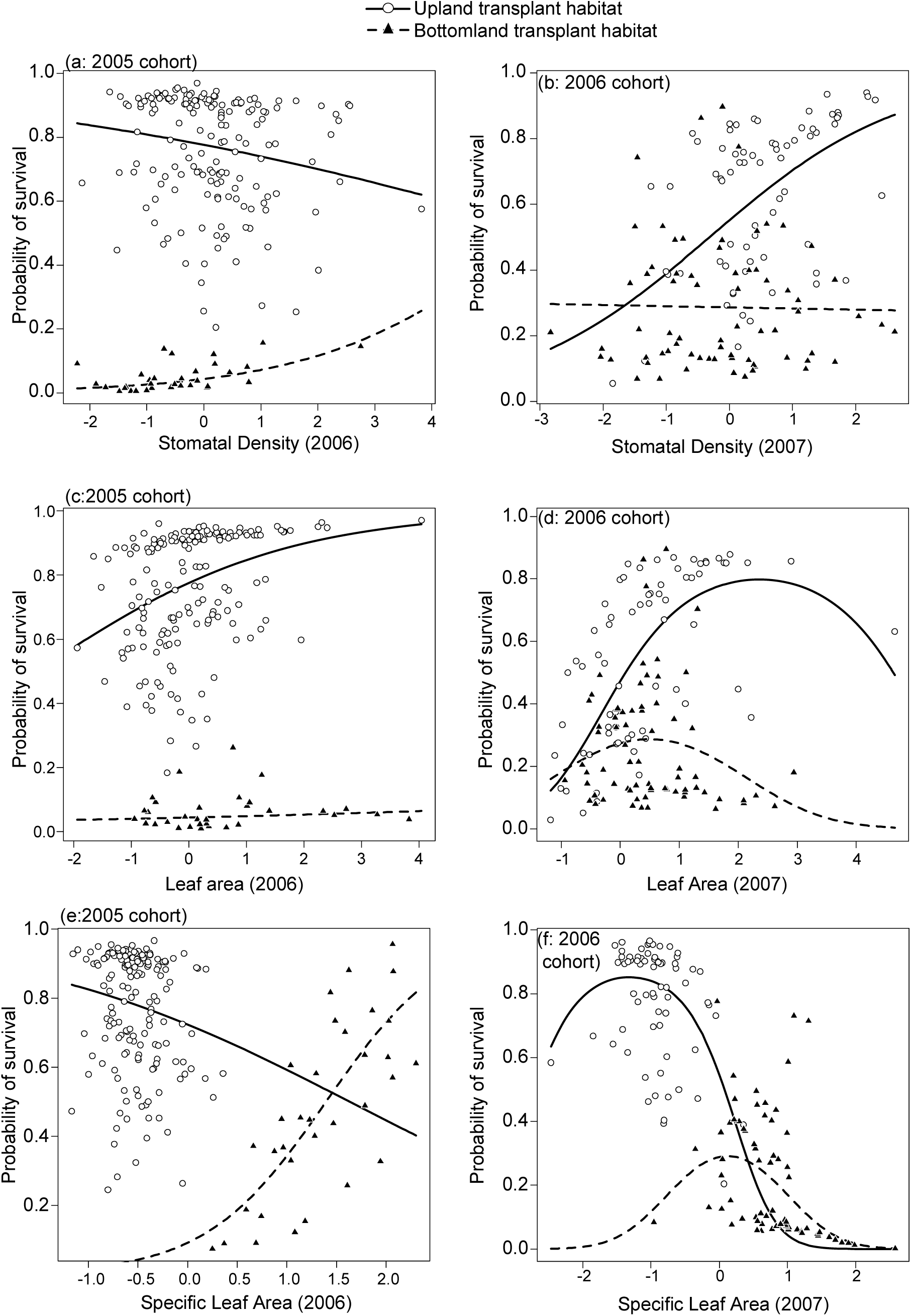
Logistic regressions evaluating viability selection on foliar traits in upland and bottomland forests for the 2005 and 2006 cohorts. For the 2005 cohort, (a)selection favored reduced stomatal density in the upland forests and increased stomatal density in the bottomlands, which contrasted with (b the 2006 cohort when selection favored greater stomatal density in the uplands with no patterns in the bottomlands. For the 2005 cohort (c), selection favored larger leaves in both sites. Quadratic selection (d) favored large leaves in the uplands in the 2006 cohort, and intermediate sized leaves in the bottomlands during the first year of trait measurement. Divergent selection favored lower specific leaf area in uplands and higher specific leaf area in the bottomlands for both cohorts (e for the 2005 cohort and f for the 2006 cohort). This divergent selection on specific leaf area was congruent with patterns of trait plasticity (Fig. 1).

